# Discovering the corky bark of *Eucalyptus suberea* from anatomical and chemical perspectives

**DOI:** 10.1101/2025.02.22.639677

**Authors:** Jorge Gominho, Ana Lourenço, Helena Patrício, Teresa Quilhó, António V. Marques

## Abstract

This paper presents the first scientific investigation of the corky bark of *Eucalyptus suberea* (cork mallee). The study involved outer bark samples from two trees with diameters of 23 cm (<u>A</u>) and 47 cm (<u>B</u>), collected from a plantation in Portugal. Anatomical analysis was performed using light and scanning electron microscopy (SEM), enabling detailed observation of bark to reveal the cork cells’ presence. The chemical characterization included summative chemical analysis, FTIR, and GC-MS/FID for suberin composition and Py-GC-MS for lignin composition. The extractive-free bark mass of *E. suberea* from trees <u>A</u> and <u>B</u> contained 29% and 25% suberin, 30% and 37% lignin, and 39% and 38% polysaccharides, respectively. The bark <u>A</u> exhibited a higher extractives content (49%) than just 14% for bark <u>B</u>. Conversely, dichloromethane extract from bark <u>A</u> was notably rich in alkanol ferulates. FTIR spectra revealed a characteristic cork tissue pattern, showing a low suberin content but high levels of polysaccharides and free carboxylic acids. The main components of *E. suberea* <u>A</u> and <u>B</u> suberins were glycerol, with 15% and 11%, ω-hydroxyacids, with 45% and 50%, and α,ω-diacids, with 30% and 17%, representing a total of ca. 90% and 78% of *E. suberea* <u>A</u> and <u>B</u> suberins mass, respectively. Both suberins showed low contents of epoxides and *vic*-diols; also, both presented the particularity of having high contents of free acid groups, especially the suberin of bark <u>B</u>, suggesting a structure with short aliphatic chains. *E. suberea* corky bark presents a lignin composition that differentiates from the cork G-lignins analyzed until now with a monomer composition exhibiting a GS character, with more G- (53-56%) than S-units (34-37%) with an S/G ratio between 0.64 to 0.66, for <u>A</u> and <u>B</u> lignins, respectively.

## 1. Introduction

*Eucalyptus suberea* Brooker & Hopper, also known as Mount Leseur mallee or cork mallee, is an endemic species on the west coast of Western Australia. A tree of the genus *Eucalyptus* and Myrtaceae that has the particularity, unlike most *Eucalyptus*, of presenting a thick grey-yellow corky bark at the base that may be flaky, thicker, and yellowish in large specimens and is smooth above (1,2). Therefore, visually and to the touch, it is similar to the virgin bark of cork oak (*Q. suber*), a characteristic of a reasonable amount of suberous tissue (3,4). This species is described as a small tree and has been found in lateritic uplands near Mount Lesueur and further inland, occurring in small populations and growing in open mallee communities, i.e., a growth habitat of eucalypt species with a height between 2 to 9m and with narrow stems; and near other *Eucalyptus* species, such as *E. lateritica*, *E. gittinsii* and *Hakea neurophylla* (1). Due to its small population, low number of plants, and unique species characteristics, it has been classified as a rare and vulnerable species under the Wildlife Conservation Act and is legally protected in Australia (5,6). The identified threats are fires, grazing, mining, invasive weeds, dieback caused by root-rot fungi, and clearing (6). In Portugal, it is located in a plantation established in 1953 within the Escaroupim National Forest, Salvaterra de Magos, Portugal. The specimens of this plantation have reasonable dimensions (20-30 m in height) and bark thicknesses that drew the authors’ attention to their possible relevance for eventual wood and cork exploitation.

While some cork-containing barks have been identified and studied, many other species remained unexplored. The first and most important bark studied is the cork from *Quercus suber* (cork oak tree), which is mainly explored to produce cork stoppers. More recently, other tree species with reasonable amounts of cork in their barks have been investigated, such as the Chinese cork oak (*Quercus variabilis*) (7), which is widely distributed throughout East Asia but only exploited in China in small quantities (8); Betula *pendula* Roth (9), *Quercus variabilis* Blume (7), *Quercus cerris* Var. cerris (10), *Pseudotsuga menziesii* (Mir.) Franco (11) and *Plathymenia reticulata* Benth. (12). Interest in these barks is steadily increasing, driven by their chemical components with diverse applications and the growing need to replace fossil-based products with sustainable bioproducts.

Cork is part of the protective tissues of plants. Therefore, it differs from wood in three aspects: its honeycomb cellular structure (7,10–15), its hydrophobic biopolymeric composition, and the assembly of polymers in the cell wall matrix (13,16–19). As wood, cork has cellulose, hemicelluloses, and lignin in its structural composition (13,18,20). Still, unlike wood, where cellulose is the main component, cork is distinguished by the presence of a 4^th^ polymeric component, the suberin, whose content varies in the cork’s tissues of different species. In *Q. suber* cork, suberin can range on average from 50% to 60% of the mass (free of extractives, 7,9,11,12,16,20,21), while in other species, lower values have been reported (25-35%, 12,22,23). Suberin has a polyester-like structure consisting mainly of poly-functional long-chain fatty α,ω-diacids, ω-hydroxyacids, and glycerol (16,19,24). Cork presents an inborn *sui generis* set of properties (13,14) that allows its use as a raw material in a wide range of industrial applications, from the well-known cork-stoppers essential for the wine industry to cutting-edge technological applications in the aerospace, automotive, construction, and other sectors (25–29). Despite being part of a diverse range of protective tissues, the commercial exploitation of cork is restricted to tree barks due to quantitative availability, productivity, and access imperatives, with the most significant production originating from cork oak forests in the Mediterranean Basin. Portugal is the major supplier of industrial cork for almost all applications, particularly those requiring better quality. The increase in consumption of cork products associated with increased diversification of applications, particularly in-home decorative products, may pose some difficulties to the cork-processing sector due to limitations in forestry production. Furthermore, the future sustainability of cork production could be compromised by climate changes associated with global warming (30,31). Despite its resilience to droughts and adaptability to poor soils (32,33), there is no absolute certainty about the cork oak species’ resistance to continued climate change over long periods. Considering these ideas, other species should also be considered for the commercial exploitation of cork. They should be the research focus, particularly in light of potential future challenges, such as using raw material species from other industries, like silver birch and Douglas-fir, or those more resilient to environmental and climate changes.

*Eucaliptus suberea* is part of a group of plants that have a reasonable amount of cork in their barks. The fact that it is a species from an area that has demanding climatic situations makes this species interesting under the scope of biorefineries. To our knowledge, no scientific study has been published regarding the bark of this species, making the anatomical and chemical examination of the cork from this *Eucalyptus* highly relevant. This work aims to provide insight into this topic material.

## 2. Material and methods

### 2.1. Sample collection and preparation

Bark samples of *Eucalyptus suberea* were collected from five trees (age unknown) present in a sown lot plantation in the Escaroupim National Forest (39°05’01.4“N, 8°43’44.3”W), Salvaterra de Magos, Portugal (Figure 2a). Tree diameters varied between 23 cm and 47 cm. Bark pieces were collected at dbh (1.30 m above ground) using a cutlass and then removed from the trunk by tapping with a hammer. The barks of the two trees with the lowest (sample <u>A</u>) and the largest diameters (sample <u>B</u>) were selected for this study. Samples were milled in a Retsch 2000 knife mill and sieved in a Retsch AS 200 apparatus with the standard mesh sieves: 10 (2 mm), 18 (1 mm), 20 (0.841 mm), 40 (0.42 mm), 60 (0.25 mm) and 80 (0.177 mm). No primary selection of bark tissue was done. The collected pieces of bark were ground as they were and milled/sieved in two stages: 1) grounding in the mill with 5x5 mm grid followed by sieving, 2) particles with more than 2 mm were ground again in the mill with 2x2 mm grid followed by sieving. The set of the three most significant fractions, 0.42-0,84 mm, 0.84-1 mm, and 1-2 mm, representing the purest suberous tissue (34) and representing more than 80% of the mass, were selected for analysis.

### 2.2. Anatomy characterization

<u>A</u> and <u>B</u> bark samples were treated with DP1500 polyethylene glycol as described in (35). Longitudinal microscopic sections approximately 17 μm thick were prepared using a Leica SM 2400 microtome and stained with chrysodine/astra blue. Sudan 4 was employed for the selective staining of suberin. Individual specimens underwent maceration with Franklin solution and were stained with astra blue. Slide preparation followed the standard procedures outlined in previous studies (36). Light microscopic observations and photomicrographs were captured using a Nikon Microphot-FXA. A sharp razor blade cut small samples with edges measuring approximately 3 mm. Their surfaces were examined using a scanning electron microscope (SEM) Hitachi TM 3030 Plus at 5 kV with varying magnifications, and the images were recorded in digital format. The bark terminology complied with the IAWA List of Microscopic Features of Bark (37).

### 2.3. Chemical characterization

The chemical summative analysis involved quantifying the ash, extractives (in dichloromethane, ethanol, and water), suberin, lignin (Klason and soluble lignin), and sugar composition. The ash content was determined by combusting 1.5 g at 525 °C overnight and weighing the residue (TAPPI standard T211 om- 93). Extractives content was assessed using a Soxhlet apparatus with a sequence of dichloromethane (6 hours), ethanol (16 hours), and water (16 hours). The difference in solid mass before and after extraction determined the extractive content. A sample of dichloromethane extract was chosen for further GC-MS analysis. The samples were oven-dried at 80 °C and then subjected to vacuum over P_2_O_5_.

Suberin depolymerization and monomeric composition analysis were conducted following the procedure outlined by Marques and Pereira (16). Approximately 200 mg of bark was depolymerized using a 0.52% sodium methoxide solution in anhydrous methanol. Two tests for suberin determination were performed for each sample size, resulting in 12 trials (six for sample <u>A</u> and six for sample <u>B</u>). Suberin analysis was carried out on all six samples (2 trees x 3 size fractions) of *E. suberea* bark. The suberin was quantified through gravimetric analysis, relying on the mass difference between the initial sample and the oven-dried (o.d.) residue mass following suberin depolymerization. The residues were used for Klason and soluble lignin determination according to TAPPI standards T222 om-88 and UM250 om-83, as well as for sugar composition analysis. Neutral monosaccharides and galacturonic acid were separated using high-pressure ion-exchange chromatography on a Dionex ICS3000 equipped with a PAD detector, employing a Carbopac PA10 (4 × 250 mm) column. The eluent consisted of NaOH + CH_3_COONa, with a 1 mL/min flow rate at 25 °C. Acetic acid was separated in a Waters 600 and measured with a UV/Vis detector at 210 nm using a Biorad Aminex 87H HPX column (300×7.8 mm), with an eluent of 10 mN H_2_SO_4_ at a flow rate of 0.6 mL/min at 30 °C.

### 2.4. FTIR analysis

Full extracted and oven dried 20-40 mesh *E. suberea* bark samples were ground with a Retsch MM200 mixer mill for 30 min. FTIR was recorded with a Bruker Vertex 70 using the standard KBr transmission technique with a resolution of 4 cm^-1^ and 32 scans accumulation. All spectra were normalized for the highest band in the 500–4000 cm^-1^ range. A private database spectrum of *Q. suber* cork was used for comparison.

### 2.5. GC-MS/FID analysis

For the GC analysis of depolymerized suberin samples and CH_2_Cl_2_ extract samples, a ThermoTrace Ultra Polaris Ion Trap apparatus from Thermo Finnigan (Austin, TX, USA) was utilized. It was equipped with a manual split/splitless injector and a fused-silica capillary column ZB-35HT (30 m × 0.25 mm × 0.10 μm) from Phenomenex (Torrance, CA, USA). GC-MS runs identified analytes, while GC-FID runs quantified them, except for the CH_2_Cl_2_ extract, where GC-MS was employed for both purposes. Thermo Excalibur software, along with the NIST, National Institute of Standards and Technology, mass spectral search program for the NIST/EPA/NIH Mass Spectral Library version 2.0a from September 2001, was used together with Wiley 6 and private spectra collection libraries for analyte identification. Peaks were integrated manually.

The GC-FID/MS analysis of suberin proceeded using the following apparatus parameters: 2 μL injection in split mode (1:30), 280°C for the injector and FID temperatures, 0.9 mL/min of helium carrier gas, and a GC-oven program temperature of 50°C (held for 2 min), followed by a ramp of 10°C/min to 210°C, 7°C/min to 280°C, and 10°C/min to 340°C (held for 6 min).

The GC-MS analysis of CH_2_Cl_2_ extractives employed the following parameters: 1 μL injection in split mode (1:20), 280°C for the injector, 0.9 mL/min of helium carrier gas, and a GC-oven program temperature of 50°C (held for 2 min), followed by ramps of 10°C/min to 130°C, 5°C/min to 200°C, 2.5°C/min to 240°C, 4°C/min to 300°C, and 5°C/min to 350°C (held for 5 min). The Polaris MS detector’s electron impact ionization energy was set to 70 eV, with an ion source temperature of 230°C and 0.3 mL/min of damping helium gas.

Each identified suberin monomer was quantified by (i) area percentage relative to the total area of all identified compounds and (ii) mass percentage of suberin considering a response factor equal to one relative to the respective internal standard: octadecanoic acid methyl ester (IS2, C19:0 OMe) for fatty acids and fatty alcohols, and butan-1,2,4-triol for glycerol. The relative response factors determined by Marques and Pereira (16) were not used because the GC/FID methodology differed. For comparison purposes with *Q. suber* cork, raw data from Marques and Pereira (16) was used.

### 2.6. Lignin composition by Py-GC-MS

Lignin composition was analyzed in the post-methanolysis residues using Py-GC-MS. Approximately 110 µg of dried bark was weighed in a quartz boat and pyrolyzed at 550°C for 1 minute in a 5150 CDS apparatus connected to an Agilent gas chromatograph (GC 7890B) equipped with a mass detector (5977B) operating at 70 eV electron impact voltage. The volatiles were separated in a fused capillary column ZB-1701 with the following specifications: 60 m x 0.25 mm i.d. x 0.25 µm film thickness. Helium was used as the carrier gas with a 1 mL/min flow. The temperatures applied were 280°C (interface) and 270°C (injector). The oven temperature program started at 40°C for 4 minutes, increased to 70°C at a rate of 10°C per minute, then to 100°C at 5°C per minute, to 265°C at 3°C per minute (held for 3 minutes), and finally to 270°C at 5°C per minute (held for 9 minutes). Compounds derived from the barks were identified by comparing the NIST2014 and Wiley computer libraries and relevant literature (38,39). The percentage of each compound was calculated based on the total area of the chromatogram. The carbohydrate-derived compounds were summed and presented as TC, while the lignin-derived compounds were denoted as TL. The percentages of *p*-hydroxyphenyl (H), guaiacyl (G), and syringyl (S) lignin-derived products were summed separately, with values reported based on total lignin. The S/G and C/L ratios were also calculated, and the H:G:S ratio was presented.

## 3. Results and discussion

### 3.1. Sample collection, grinding, and selection

The granulate of *E. suberea* exhibited a collection of large, rounded, soft, and lightweight granules resembling *Q. suber* cork granulate (Fig 1.a). In Fig 1.b, the mass distribution is presented after sieving. This milling/sieving result agrees with that observed for a material whose behavior in a knife mill corresponds to a cork, which is poor in hard and brittle material (9,22). After sieving, the three largest particle size fractions represented 86% and 83% of the <u>A</u> and <u>B</u> mass samples. Sample <u>B</u> milling resulted in a 3% higher fines content derived from the hole’s harder tissues (Fig 2.c,d,e).

**Fig 1.**
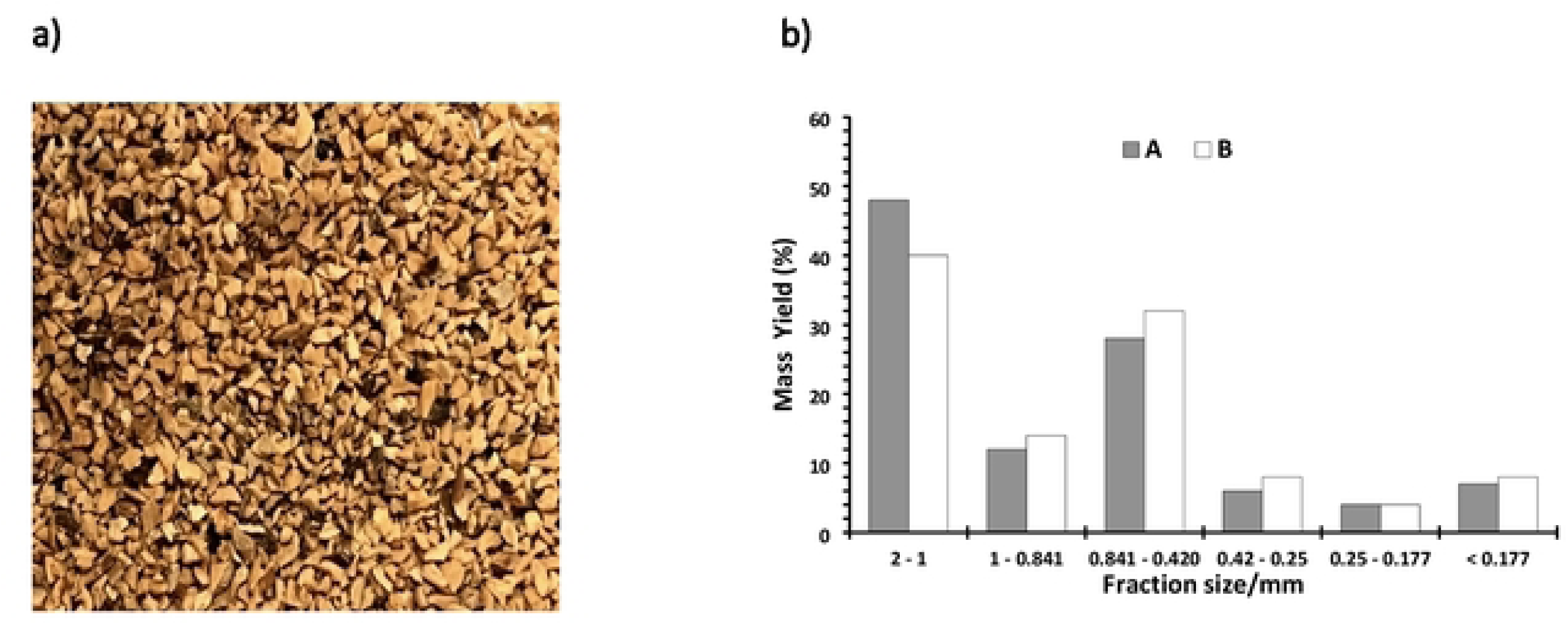
The *E. suberea* bark: a) image of the granulate; b) the mass yield distribution (%) of sieved fractions.

**Fig 2.**
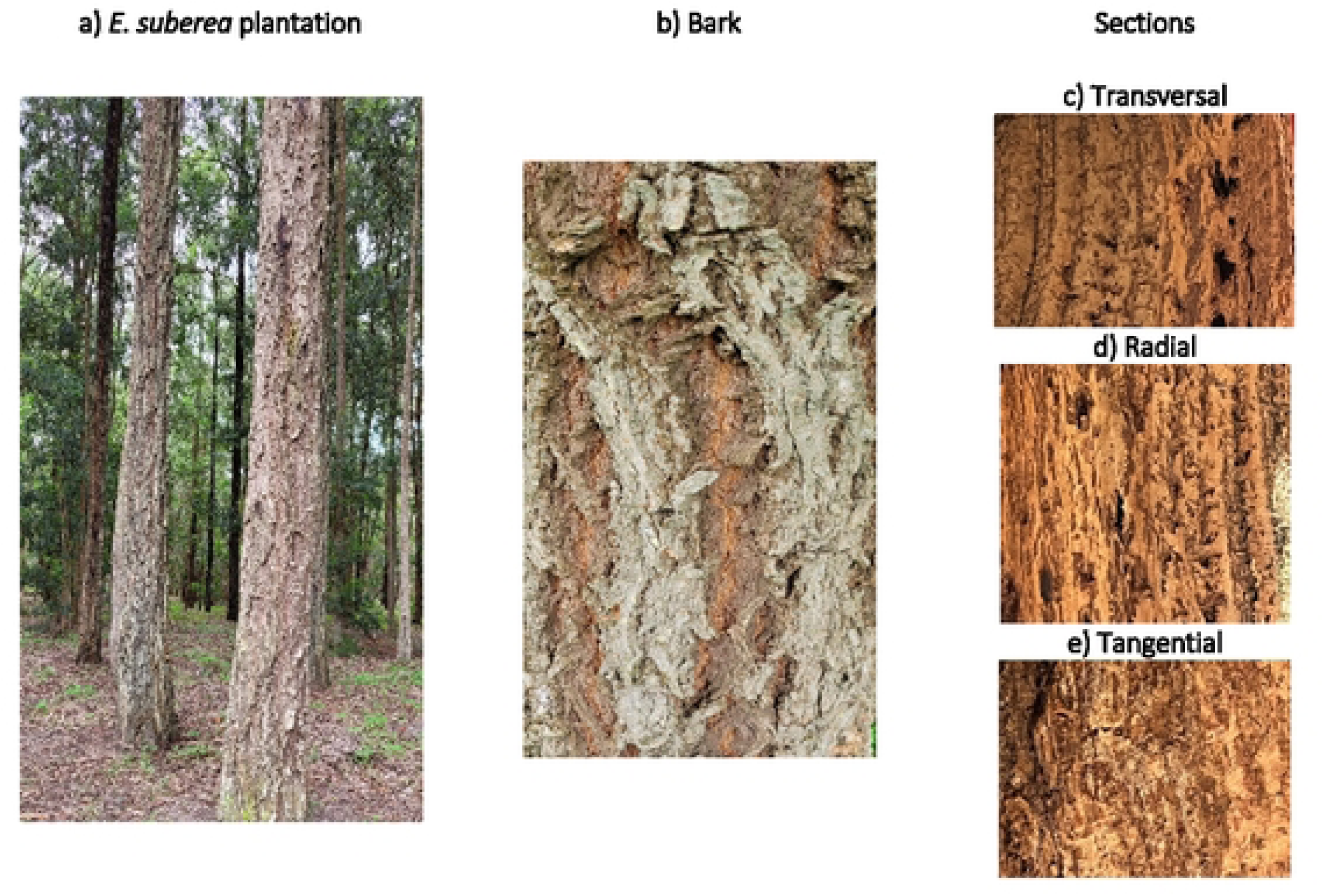
a) General overview of the *E. suberea* plantation in Escaroupim National Forest; b) close-up of the bark; c-e) detailed sections of *E. suberea* bark.

Unlike what happens with other corky barks, in which the cork is in a striped mixture of layers, or pockets, with secondary phloem, such as in Douglas-fir (11,40) and *Q. cerris* (10,23), or the cork is stratified into layers that separate, reducing the possibility of obtaining coarse granules, such as in Birch (9,22). *E. suberea* bark did not require any physical separation between phellem (cork) and phloem tissues; that is, the entire bark could be crushed as it is, with a high yield of coarse granules with granulometric dimensions that depend on the sieves used in the mill.

### 3.2. Anatomy characterization

The bark of *E. suberea* is rough, presenting wide and deep growth cracks in the axial direction. The surface color is quite variable, from a more grayish color similar to the virgin cork of *Q. suber* to a darker color (Fig 2.a and b) as also mentioned in the literature (1,2). In the transversal (Fig 2.c) and radial sections (Fig 2.d), it is possible to see the annual rings and the holes originated from the growth stress.

Bark consists of all tissues outside the vascular cambium, including the secondary phloem and the rhytidome (37), resulting in a highly complex and heterogeneous structure. The bark of the *Eucalyptus* has been examined, and the anatomical characterization of various species is documented in the literature (41–46). In this study, only the rhytidome was analyzed. The rhytidome encased the periderms (comprising phelloderm, phellogen, and phellem-cork cells) and dead secondary phloem tissue between them. The analysis of the rhytidome was conducted using scanning electron microscopy and light microscopy, enabling the observation of numerous cork cells (Fig 3 A-C; Fig 4 A-B), secretory structures (Fig 3.C), and an increased accumulation of extractives in the sclerified cells (Fig 4.C).

**Fig 3.**
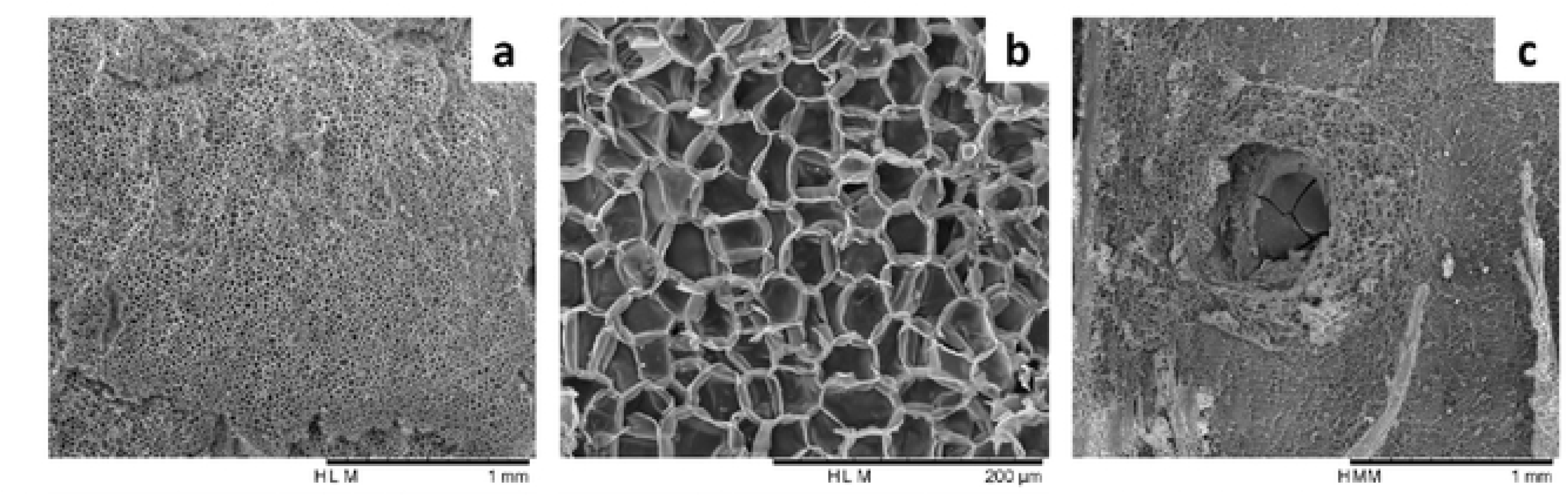
Rhytidome of the *Eucalyptus suberea* (tangential section) visualized by SEM. **a-b)** Cork cells **c)** secretory structure (gland).

**Fig 4.**
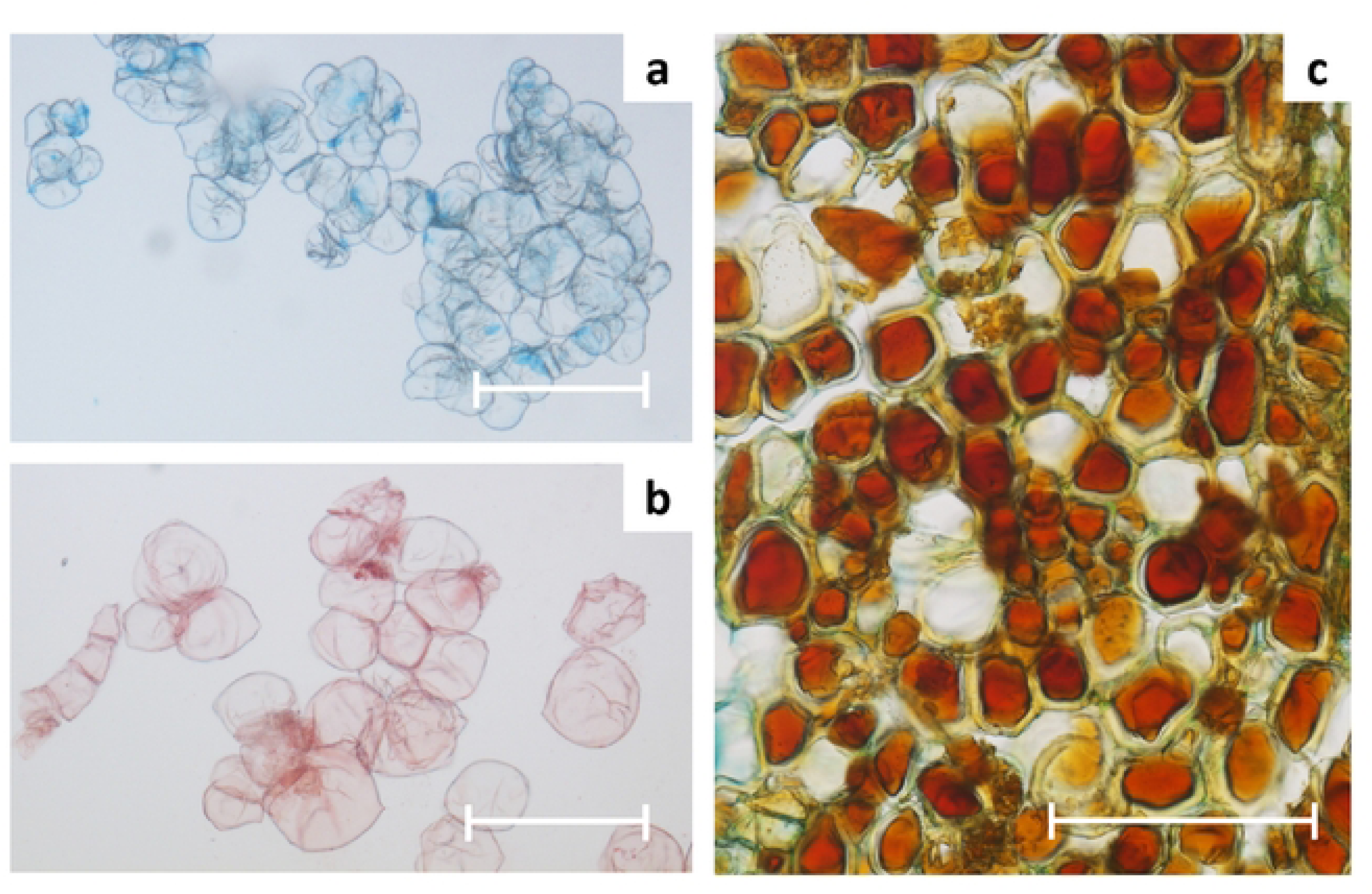
Cells of the *Eucalyptus subereo* rhytidome. **a)** Suberized cells are stained with Astra blue. **b)** Suberized cells are stained with Sudan IV. c) lignified cells with abundant extractives (radial section). Scale bar: a, c = 150 µm, b = 100 µm.

The presence of abundant cork cells in *E. suberea* makes it different from other species of eucalypts. The phellem cells in the periderm of *Eucalyptus* are, in general, poor suberized, *e.g.*, in *E. globulus* (43) in opposition to other species such as *Q. cerris* (47), *Pseudotsuga menziesii* (36), and *Plathymenia reticulata* (12) with a large portion of cork, or *Q. suber* (13,48) and *Q. variabilis* (49), which have a periderm with a thick cork layer. The large amount of cork in the rhytidome of *E. suberea* reflects its unmistakable singular external appearance (corky bark).

### 3.3. FTIR analysis

Fig 5.a presents the FTIR spectra of *E. suberea* samples compared to those of *Q. suber* cork (the spectra have been normalized to the highest peaks). The spectra of both *E. suberea* barks <u>A</u> and <u>B</u>, apart from the differences in the carbonyl bands, are pretty similar and exhibit the typical cork tissue pattern, indicating suberin’s presence. This is characterized by C-H stretch bands between 2800 and 3000 cm⁻¹, which correspond to the long aliphatic chains of suberinic acids and the strong stretch vibration bands of fatty esters. The carbonyl C=O stretch occurs at 1740 cm⁻¹ and 1736 cm⁻¹ for *E. suberea* <u>A</u> and <u>B</u>, respectively, and at 1742 cm⁻¹ for cork oak, alongside the asymmetric ester C-C(=O)-O stretch at 1160 cm⁻¹ and 1161 cm⁻¹. The relative intensity of the C=O bands of both *E. suberea* samples is weaker than that of the C=O band from *Q. suber*. In contrast, the O-H stretch bands of *E. suberea* are higher and broader than the OH band in *Q. suber* cork, as are the C-OH stretch bands at 1036 cm^-1^ and 1037 cm^-1^. These differences indicate a lower suberin content in the *E. suberea* barks than in *Q. suber* and, conversely, a higher content of carbohydrates, which will be confirmed by chemical analysis.

**Fig 5.**
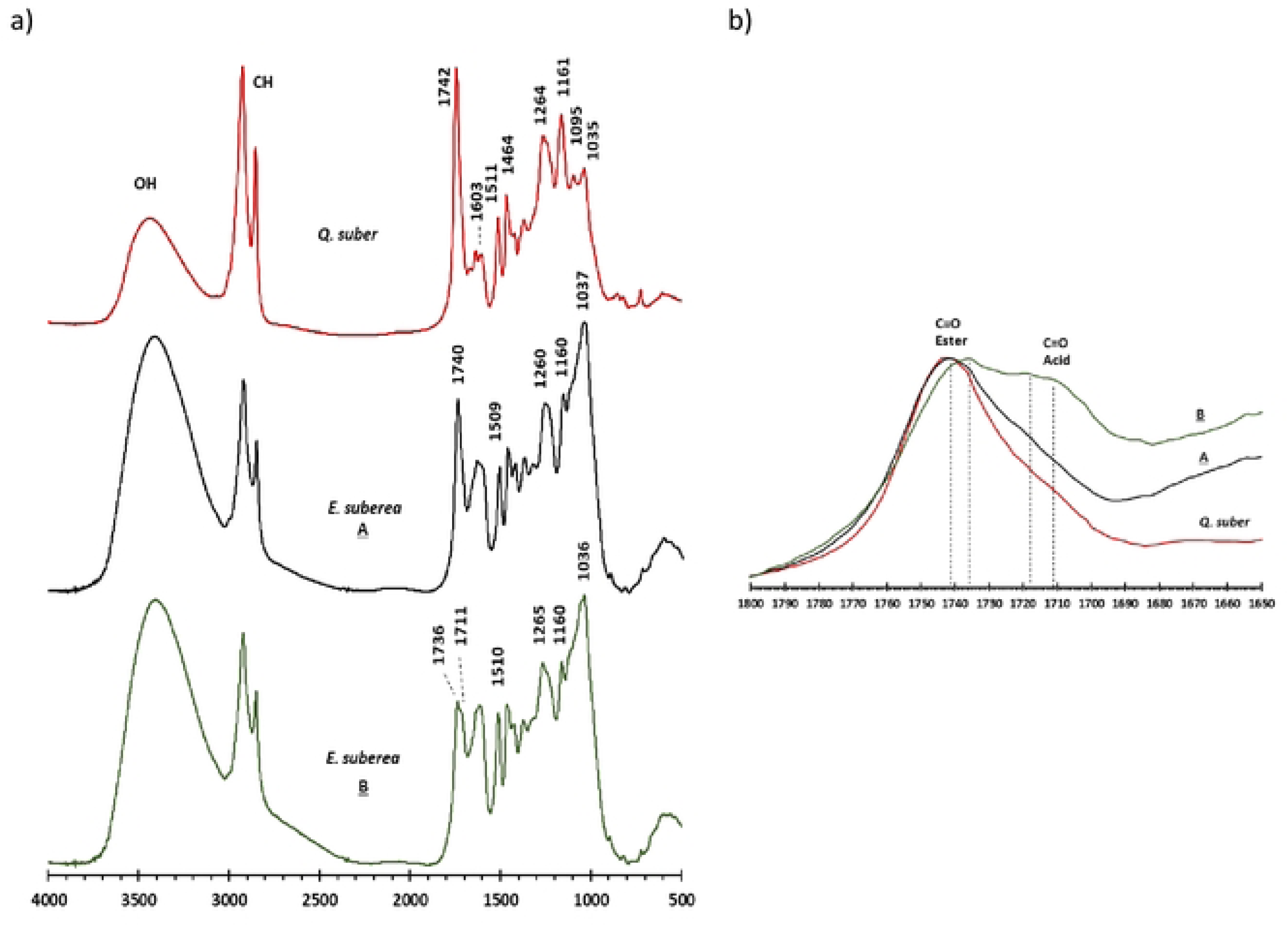
FTIR spectra of *E. suberea* bark samples <u>A</u>, <u>B</u>, and *Q. suber* cork **(a),** and the corresponding C=O stretch bands at around1740 and between 1720-1730 cm^−1^ (b).

In line with the broader width of the O-H band in the *Q. suberea* samples, which indicates the presence of COOH groups, it is also observed in the case of *E. suberea*, particularly in sample <u>B</u>, that the C=O band is less distinct, displaying two frequencies: one corresponding to the C=O of ester groups at 1736 cm^-1^ and the other at 1711 cm^-1^ corresponding to carboxylic acid. This suggests that *E. suberea* suberin has a higher content of free acid groups than cork oak suberin (16), which is more significant in sample <u>B</u>. Furthermore, it is noted that the free acid groups are situated in varying structural environments, resulting in the presence of the two maxima at 1711 cm^-1^ and 1718 cm^-1^ (Figure 5.b). Suberin analysis via GC/FID will confirm this result.

The lignin in *E. suberea* barks is indicated by 1509 cm^-1^ and 1510 cm^-1^ bands from aromatic skeletal vibrations. The frequency of these bands decreases slightly for lignins with a higher content of syringyl rings; therefore, these samples are expected to have lignin with a composition containing more syringyl rings than cork oak or Douglas-fir corks (17,50,51). A more careful observation of the area between 900 cm^-1^ and 1200 cm^-1^ allows us to verify that the spectra of *E. suberea* exhibit the presence of a band at 1128 cm^-1^ that is a characteristic of GS lignins (51), while this band is missing in *Q. suber* cork.

### 3.4. Chemical composition

The chemical summative analysis for inorganics, extractable, and structural components of *E. suberea* bark are presented in Table 1. Overall, it is possible to see some chemical differences between the two barks. Suberin content of *E. suberea* bark was higher in sample <u>A</u> (28.8%) compared to sample <u>B</u> (24.6% of extractive free o.d. material) and being *ca.* half that found in cork oaks (16,20,21) or Douglas-fir (21,40). This result is following FTIR spectra analysis. Even though the content and composition of suberin can vary greatly with the sample origin and with the methodology used in its determination (52), when we compare the yields of suberin in *E. suberea* bark with other species, we find that the value is in the lower fringe of the range observed in the literature for corky barks with relevant cork amounts, namely, *Plathymenia reticulata* with 28.3% (12), *Q. cerris* with 34.2% (23), *Quercus variabilis* with 41.8% (7), *Quercus suber* with 51.1% (20) and *Betula pendula* with 53.4% (9).

**Table 1.** Summative analysis of *E. suberea* bark samples and *Q. suber* cork as reference.

|  | Oven-dried |  | Oven-dried without extractives |  | *Q. suber*<sup>a</sup> |
| --- | --- | --- | --- | --- | --- |
|  | <u>A</u> | <u>B</u> | <u>A</u> | <u>B</u> |  |
| Ash | 2.4 | 0.5 | 2.8 | 0.9 | - |
| Extractives | 14.2 | 49.4 | - | - | 16.2 |
| Dichloromethane | 4.6 | 11.4 | - | - | 5.8 |
| Ethanol | 3.9 | 16.6 | - | - | 5.9 |
| Water | 5.8 | 21.4 | - | - | 4.5 |
| Suberin | 24.7 | 12.4 | 28.8 | 24.6 | 42.8 |
| Lignin | 25.6 | 18.6 | 29.8 | 36.7 | 22.0 |
| Klason | 25.5 | 18.5 | 29.7 | 36.6 | 21.1 |
| Soluble | 0.09 | 0.05 | 0.1 | 0.1 | 0.9 |
| Polysaccharides | 33.1 | 19.1 | 38.6 | 37.8 | 19.0 |
| Rhamnose <sup>b</sup> | 1.4<sup>b</sup> | 1.1<sup>b</sup> | - | - | 0.5 |
| Arabinose | 13.1 | 6.7 | - | - | 18.0 |
| Xylose | 28.5 | 27.4 | - | - | 25.1 |
| Glucose | 47.0 | 54.8 | - | - | 46.1 |
| Galactose | 5.6 | 5.5 | - | - | 7.3 |
| Mannose | 1.7 | 2.0 | - | - | 3.0 |
| Galacturonic acid | 2.1 | 2.0 | - | - | - |
| Glucuronic acid | 0.4 | 0.4 | - | - | - |
| Acetic acid | 0.1 | 0.0 | - | - | - |

<sup>a</sup> Data from H. Pereira 2013 (20). <sup>b</sup> Monosaccharides relative composition to polysaccharides.

The biggest difference between the *E. suberea* samples <u>A</u> and <u>B</u> relates to the extractive content, with the larger tree (bark <u>B</u>) exhibiting an extractive level in its bark that is 3.5 times higher. The exact reason is unknown but may be associated with defense mechanisms linked to a potentially longer lifetime of environmental exposure. The small tree (bark <u>A</u>) shows an extractive content at the lower end of the range reported in the literature for other species, comparable to *Q. suber*, *Q. cerris*, and *P. reticulata* (12,20,23). In contrast, the bark of larger trees surpasses the highest published values, reaching 37.5% for Douglas-fir cork (21). Lignin and polysaccharide values are within the range observed for other species. The monosaccharide composition aligns with that typically seen in this type of material, with glucose making up the largest proportion at around 50%, followed by xylose and then arabinose, with only trace amounts of mannose and rhamnose, similar to what is observed in others corks. In the case of *Q. suber,* polysaccharides represent ca. 20% of extractive-free cork, where glucose and xylose represent roughly 50% and 35% of all the sugars liberated by total hydrolysis. All the other pentoses present, such as arabinose, mannose and rhamnose, represent *ca*. 10% (13,20).

### 3.5. Extractives lipophilic composition

Table 2 presents the composition of lipophilic extractives of *E. suberea* barks. The identified compounds account for 85% and 87% of the GC-MS peak areas for samples <u>A</u> and <u>B</u>, respectively. Bark from the largest tree contains 2.5 times more lipophilic extract than the smallest tree, and this increase is attributed to a significant rise in fatty components, n-alkanols, acids, monoglycerides, and ferulates of n-alkanols. Ferulates and α,ω-bifunctional acids comprise 27% and 25% of this composition extract.

**Table 2.**
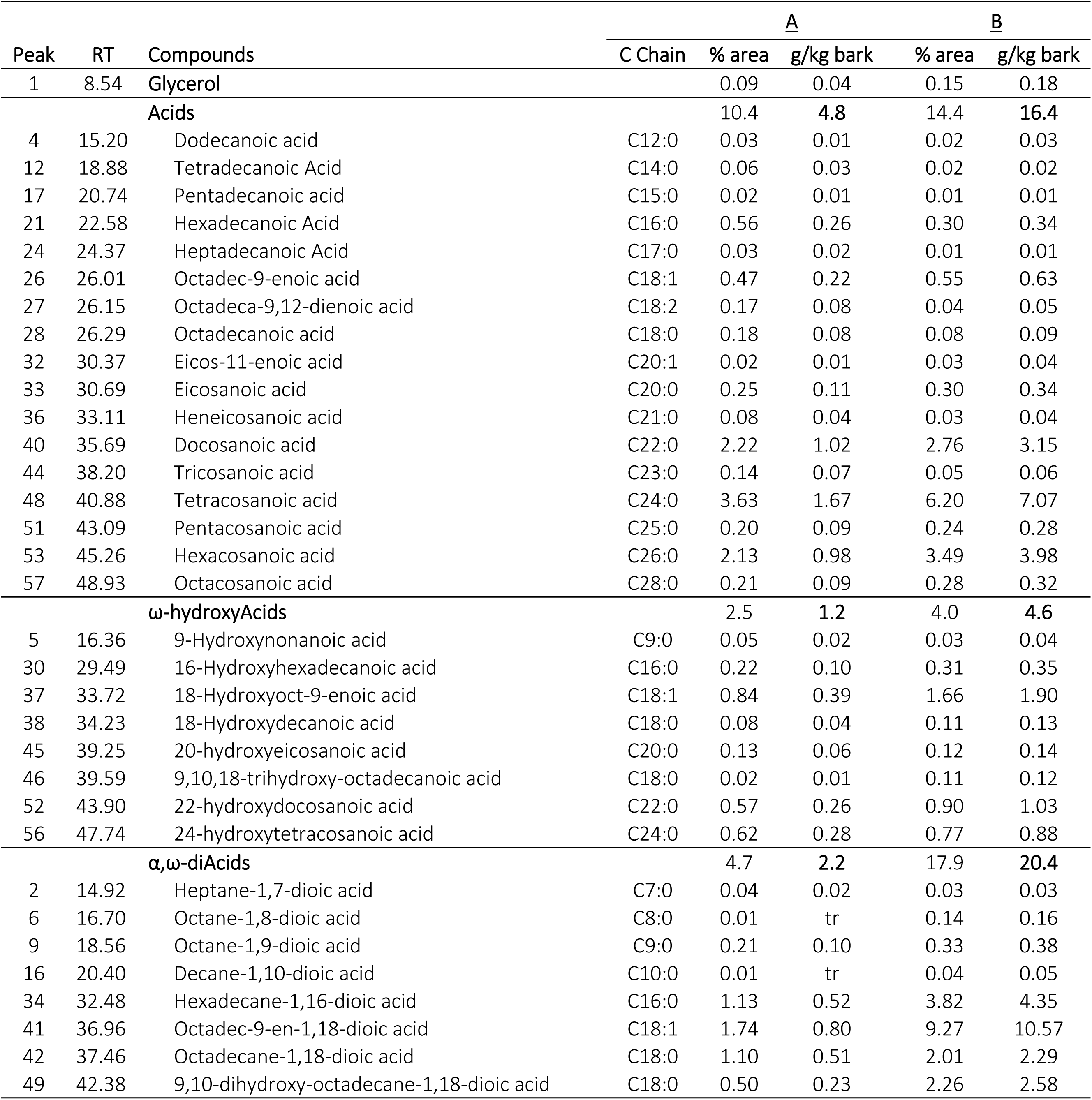
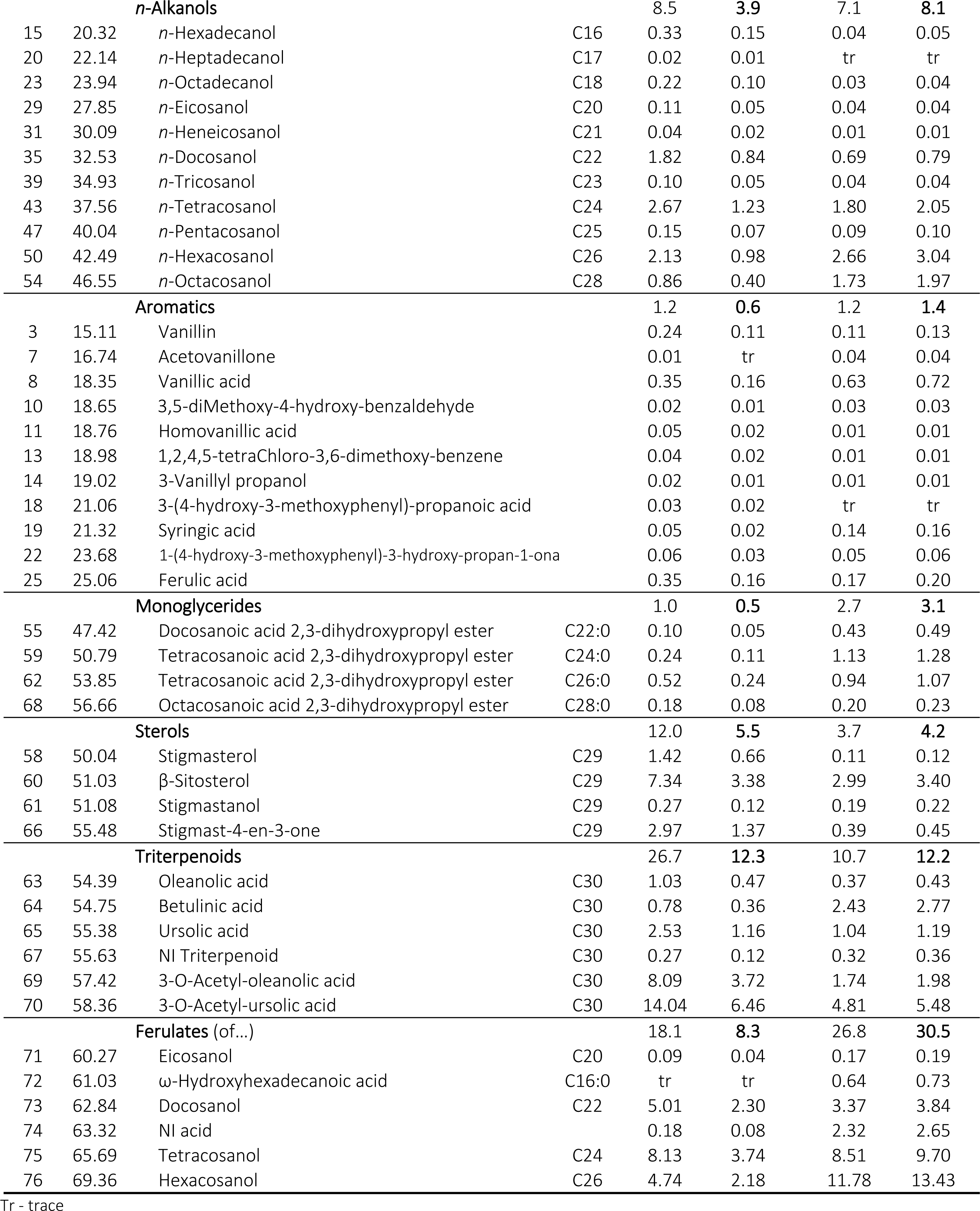
Composition of dichloromethane extract from *E. suberea* barks. Results are presented in the GC-MS relative area and mass percentage relative to o.d. bark, considering that all the area represents the extracted mass.

| Peak | RT | Compounds | C Chain | A |  | B |  |
| --- | --- | --- | --- | --- | --- | --- | --- |
|  |  |  |  | % area | g/kg bark | % area | g/kg bark |
| 1 | 8.54 | Glycerol |  | 0.09 | 0.04 | 0.15 | 0.18 |
|  |  | **Acids** |  | 10.4 | **4.8** | 14.4 | **16.4** |
| 4 | 15.20 | Dodecanoic acid | C12:0 | 0.03 | 0.01 | 0.02 | 0.03 |
| 12 | 18.88 | Tetradecanoic Acid | C14:0 | 0.06 | 0.03 | 0.02 | 0.02 |
| 17 | 20.74 | Pentadecanoic acid | C15:0 | 0.02 | 0.01 | 0.01 | 0.01 |
| 21 | 22.58 | Hexadecanoic Acid | C16:0 | 0.56 | 0.26 | 0.30 | 0.34 |
| 24 | 24.37 | Heptadecanoic Acid | C17:0 | 0.03 | 0.02 | 0.01 | 0.01 |
| 26 | 26.01 | Octadec-9-enoic acid | C18:1 | 0.47 | 0.22 | 0.55 | 0.63 |
| 27 | 26.15 | Octadeca-9,12-dienoic acid | C18:2 | 0.17 | 0.08 | 0.04 | 0.05 |
| 28 | 26.29 | Octadecanoic acid | C18:0 | 0.18 | 0.08 | 0.08 | 0.09 |
| 32 | 30.37 | Eicos-11-enoic acid | C20:1 | 0.02 | 0.01 | 0.03 | 0.04 |
| 33 | 30.69 | Eicosanoic acid | C20:0 | 0.25 | 0.11 | 0.30 | 0.34 |
| 36 | 33.11 | Heneicosanoic acid | C21:0 | 0.08 | 0.04 | 0.03 | 0.04 |
| 40 | 35.69 | Docosanoic acid | C22:0 | 2.22 | 1.02 | 2.76 | 3.15 |
| 44 | 38.20 | Tricosanoic acid | C23:0 | 0.14 | 0.07 | 0.05 | 0.06 |
| 48 | 40.88 | Tetracosanoic acid | C24:0 | 3.63 | 1.67 | 6.20 | 7.07 |
| 51 | 43.09 | Pentacosanoic acid | C25:0 | 0.20 | 0.09 | 0.24 | 0.28 |
| 53 | 45.26 | Hexacosanoic acid | C26:0 | 2.13 | 0.98 | 3.49 | 3.98 |
| 57 | 48.93 | Octacosanoic acid | C28:0 | 0.21 | 0.09 | 0.28 | 0.32 |
|  |  | **ω-hydroxyAcids** |  | 2.5 | **1.2** | 4.0 | **4.6** |
| 5 | 16.36 | 9-Hydroxynonanoic acid | C9:0 | 0.05 | 0.02 | 0.03 | 0.04 |
| 30 | 29.49 | 16-Hydroxyhexadecanoic acid | C16:0 | 0.22 | 0.10 | 0.31 | 0.35 |
| 37 | 33.72 | 18-Hydroxyoct-9-enoic acid | C18:1 | 0.84 | 0.39 | 1.66 | 1.90 |
| 38 | 34.23 | 18-Hydroxydecanoic acid | C18:0 | 0.08 | 0.04 | 0.11 | 0.13 |
| 45 | 39.25 | 20-hydroxyeicosanoic acid | C20:0 | 0.13 | 0.06 | 0.12 | 0.14 |
| 46 | 39.59 | 9,10,18-trihydroxy-octadecanoic acid | C18:0 | 0.02 | 0.01 | 0.11 | 0.12 |
| 52 | 43.90 | 22-hydroxydocosanoic acid | C22:0 | 0.57 | 0.26 | 0.90 | 1.03 |
| 56 | 47.74 | 24-hydroxytetracosanoic acid | C24:0 | 0.62 | 0.28 | 0.77 | 0.88 |
|  |  | **$\alpha,\omega$-diAcids** |  | 4.7 | **2.2** | 17.9 | **20.4** |
| 2 | 14.92 | Heptane-1,7-dioic acid | C7:0 | 0.04 | 0.02 | 0.03 | 0.03 |
| 6 | 16.70 | Octane-1,8-dioic acid | C8:0 | 0.01 | tr | 0.14 | 0.16 |
| 9 | 18.56 | Octane-1,9-dioic acid | C9:0 | 0.21 | 0.10 | 0.33 | 0.38 |
| 16 | 20.40 | Decane-1,10-dioic acid | C10:0 | 0.01 | tr | 0.04 | 0.05 |
| 34 | 32.48 | Hexadecane-1,16-dioic acid | C16:0 | 1.13 | 0.52 | 3.82 | 4.35 |
| 41 | 36.96 | Octadec-9-en-1,18-dioic acid | C18:1 | 1.74 | 0.80 | 9.27 | 10.57 |
| 42 | 37.46 | Octadecane-1,18-dioic acid | C18:0 | 1.10 | 0.51 | 2.01 | 2.29 |
| 49 | 42.38 | 9,10-dihydroxy-octadecane-1,18-dioic acid | C18:0 | 0.50 | 0.23 | 2.26 | 2.58 |

| *n*-Alkanols |  |  |  | 8.5 | 3.9 | 7.1 | 8.1 |
| --- | --- | --- | --- | --- | --- | --- | --- |
| 15 | 20.32 | *n*-Hexadecanol | C16 | 0.33 | 0.15 | 0.04 | 0.05 |
| 20 | 22.14 | *n*-Heptadecanol | C17 | 0.02 | 0.01 | tr | tr |
| 23 | 23.94 | *n*-Octadecanol | C18 | 0.22 | 0.10 | 0.03 | 0.04 |
| 29 | 27.85 | *n*-Eicosanol | C20 | 0.11 | 0.05 | 0.04 | 0.04 |
| 31 | 30.09 | *n*-Heneicosanol | C21 | 0.04 | 0.02 | 0.01 | 0.01 |
| 35 | 32.53 | *n*-Docosanol | C22 | 1.82 | 0.84 | 0.69 | 0.79 |
| 39 | 34.93 | *n*-Tricosanol | C23 | 0.10 | 0.05 | 0.04 | 0.04 |
| 43 | 37.56 | *n*-Tetracosanol | C24 | 2.67 | 1.23 | 1.80 | 2.05 |
| 47 | 40.04 | *n*-Pentacosanol | C25 | 0.15 | 0.07 | 0.09 | 0.10 |
| 50 | 42.49 | *n*-Hexacosanol | C26 | 2.13 | 0.98 | 2.66 | 3.04 |
| 54 | 46.55 | *n*-Octacosanol | C28 | 0.86 | 0.40 | 1.73 | 1.97 |
| Aromatics |  |  |  | 1.2 | 0.6 | 1.2 | 1.4 |
| 3 | 15.11 | Vanillin |  | 0.24 | 0.11 | 0.11 | 0.13 |
| 7 | 16.74 | Acetovanillone |  | 0.01 | tr | 0.04 | 0.04 |
| 8 | 18.35 | Vanillic acid |  | 0.35 | 0.16 | 0.63 | 0.72 |
| 10 | 18.65 | 3,5-diMethoxy-4-hydroxy-benzaldehyde |  | 0.02 | 0.01 | 0.03 | 0.03 |
| 11 | 18.76 | Homovanillic acid |  | 0.05 | 0.02 | 0.01 | 0.01 |
| 13 | 18.98 | 1,2,4,5-tetraChloro-3,6-dimethoxy-benzene |  | 0.04 | 0.02 | 0.01 | 0.01 |
| 14 | 19.02 | 3-Vanillyl propanol |  | 0.02 | 0.01 | 0.01 | 0.01 |
| 18 | 21.06 | 3-(4-hydroxy-3-methoxyphenyl)-propanoic acid |  | 0.03 | 0.02 | tr | tr |
| 19 | 21.32 | Syringic acid |  | 0.05 | 0.02 | 0.14 | 0.16 |
| 22 | 23.68 | 1-(4-hydroxy-3-methoxyphenyl)-3-hydroxy-propan-1-ona |  | 0.06 | 0.03 | 0.05 | 0.06 |
| 25 | 25.06 | Ferulic acid |  | 0.35 | 0.16 | 0.17 | 0.20 |
| Monoglycerides |  |  |  | 1.0 | 0.5 | 2.7 | 3.1 |
| 55 | 47.42 | Docosanoic acid 2,3-dihydroxypropyl ester | C22:0 | 0.10 | 0.05 | 0.43 | 0.49 |
| 59 | 50.79 | Tetracosanoic acid 2,3-dihydroxypropyl ester | C24:0 | 0.24 | 0.11 | 1.13 | 1.28 |
| 62 | 53.85 | Tetracosanoic acid 2,3-dihydroxypropyl ester | C26:0 | 0.52 | 0.24 | 0.94 | 1.07 |
| 68 | 56.66 | Octacosanoic acid 2,3-dihydroxypropyl ester | C28:0 | 0.18 | 0.08 | 0.20 | 0.23 |
| Sterols |  |  |  | 12.0 | 5.5 | 3.7 | 4.2 |
| 58 | 50.04 | Stigmasterol | C29 | 1.42 | 0.66 | 0.11 | 0.12 |
| 60 | 51.03 | β-Sitosterol | C29 | 7.34 | 3.38 | 2.99 | 3.40 |
| 61 | 51.08 | Stigmastanol | C29 | 0.27 | 0.12 | 0.19 | 0.22 |
| 66 | 55.48 | Stigmast-4-en-3-one | C29 | 2.97 | 1.37 | 0.39 | 0.45 |
| Triterpenoids |  |  |  | 26.7 | 12.3 | 10.7 | 12.2 |
| 63 | 54.39 | Oleanolic acid | C30 | 1.03 | 0.47 | 0.37 | 0.43 |
| 64 | 54.75 | Betulinic acid | C30 | 0.78 | 0.36 | 2.43 | 2.77 |
| 65 | 55.38 | Ursolic acid | C30 | 2.53 | 1.16 | 1.04 | 1.19 |
| 67 | 55.63 | NI Triterpenoid | C30 | 0.27 | 0.12 | 0.32 | 0.36 |
| 69 | 57.42 | 3-O-Acetyl-oleanolic acid | C30 | 8.09 | 3.72 | 1.74 | 1.98 |
| 70 | 58.36 | 3-O-Acetyl-ursolic acid | C30 | 14.04 | 6.46 | 4.81 | 5.48 |
| Ferulates (of...) |  |  |  | 18.1 | 8.3 | 26.8 | 30.5 |
| 71 | 60.27 | Eicosanol | C20 | 0.09 | 0.04 | 0.17 | 0.19 |
| 72 | 61.03 | ω-Hydroxyhexadecanoic acid | C16:0 | tr | tr | 0.64 | 0.73 |
| 73 | 62.84 | Docosanol | C22 | 5.01 | 2.30 | 3.37 | 3.84 |
| 74 | 63.32 | NI acid |  | 0.18 | 0.08 | 2.32 | 2.65 |
| 75 | 65.69 | Tetracosanol | C24 | 8.13 | 3.74 | 8.51 | 9.70 |
| 76 | 69.36 | Hexacosanol | C26 | 4.74 | 2.18 | 11.78 | 13.43 |

Tr - trace

Despite various analytical methodologies and natural variability, the literature indicates that the typical composition pattern of these extracts is rich in triterpenes and triterpenic acids, specifically betulin, friedelin, and betulinic acid (7,9,40,53), with exceptions such as Douglas-fir, where flavonoids were identified in greater quantities (40). The lipophilic extract of *E. suberea* exhibits a different pattern, dominated by fatty components, particularly ferulates. Ferulic acid and related ester compounds are recognized for their anti-oxidant and other biological activities (54), and, as previously highlighted, the increased content of lipophilic extractives and ferulates may be linked to an environmentally protective response of the tree. As reported in numerous studies of the valorization of the plant barks, the extracts from *E. suberea* could also contribute to a sustainable source for future biorefining developments.

### 3.6. Suberin analysis

Fig 6 shows an example of a GC-FID chromatogram for *E. suberea* suberin analysis. Table 3 presents the full GC-FID results for the suberin composition, and Table 4 summarizes the results by component family.

**Fig 6.**
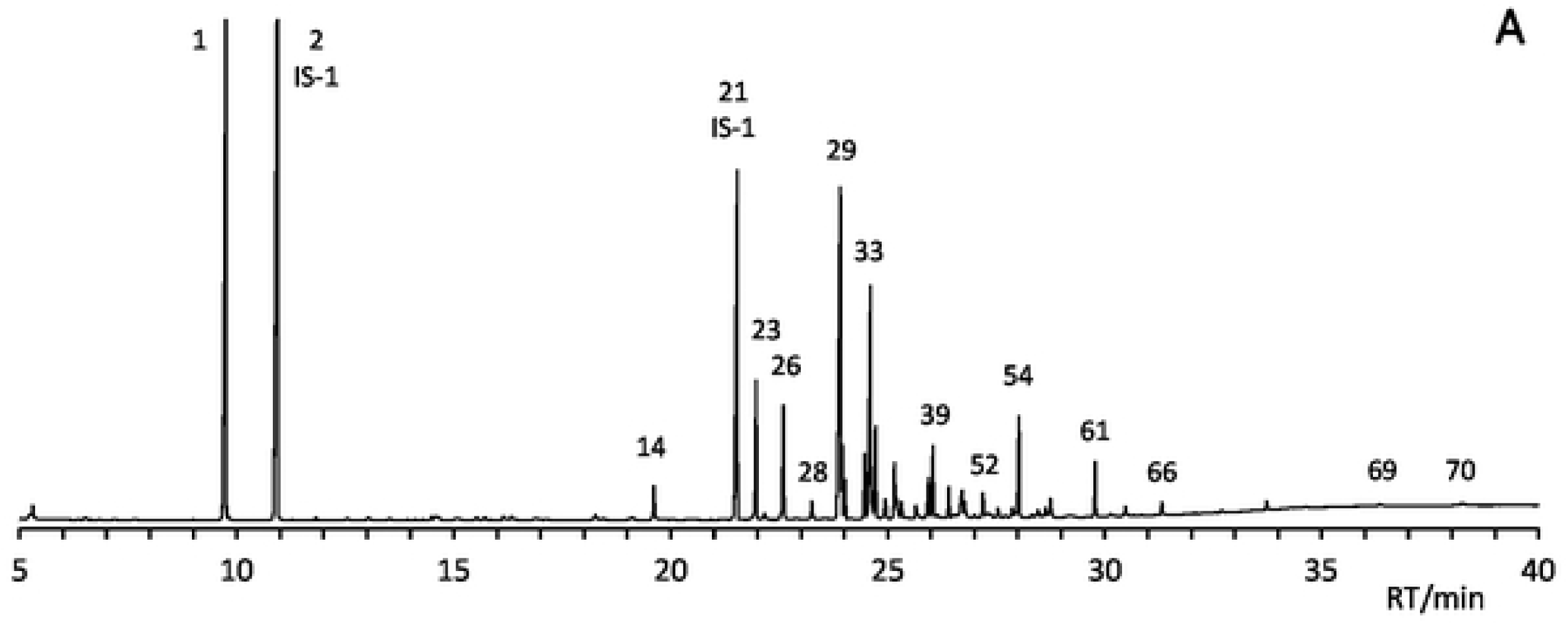
GC/FIO chromatogram example of Eucalyptus suberea suberin sample A. Numbers corresponds to identificationsin Table 3.

**Table 3.** GC-MS identification and GC-FID quantification of suberin components, by area and mass percentages, from *Eucalyptus suberea* bark and *Quercus suber* cork. Acid groups were identified and analyzed as methyl ester except when indicated.

| Peak | RT | ID <sup>a</sup> | % Area <sup>b</sup> |  |  | % Mass to Suberin <sup>c</sup> |  |  |
| --- | --- | --- | --- | --- | --- | --- | --- | --- |
|  |  |  | A | B | *Q. suber*<sup>d</sup> | A | B | *Q. suber*<sup>d</sup> |
| 1 | 9.7 | Glycerol | 19.45 | 12.48 | 23.59 | 14.91 | 10.94 | 17.27 |
| 2 | 10.9 | Butan-1,2,4-triol (IS-1) | 24.98 | 32.62 | 18.29 |  |  |  |
| 3 | 14.9 | ω-Hydroxy acid C9:0 | 0.02 | 0.09 | 0.02 | 0.03 | 0.19 | 0.02 |
| 4 | 15.1 | Vanillin | 0.06 | 0.11 | 0.06 |  |  |  |
| 5 | 15.7 | ω-Hydroxy acid C9:0 TMS ester | 0.06 | 0.17 | | 0.12 | 0.38 | |
| 6 | 16.2 | $\alpha,\omega$-Diacid C9:0 α-TMS ester | 0.11 | 0.12 | 0.05 | 0.22 | 0.27 | 0.07 |
| 7 | 16.2 | 4-Hydroxy-3-methoxy-benzoic acid | 0.06 | 0.06 |  | 0.11 | 0.13 |  |
| 8 | 16.9 | Vanillic acid TMS ester | 0.07 | 0.08 | 0.03 |  |  |  |
| 9 | 17.0 | 3,4-Dihydroxybenzoic acid TMS ester | 0.01 | 0.04 | 0.02 |  |  |  |
| 10 | 18.3 | *cis*-Ferulic acid | 0.18 | 0.16 | 0.01 |  |  |  |
| 11 | 18.4 | Acid C16:0 | 0.02 | 0.02 |  | 0.04 | 0.05 |  |
| 12 | 19.2 | Acid C16:0 TMS ester | 0.06 | 0.05 |  | 0.11 | 0.10 |  |
| 13 | 19.5 | 3,4-Dihydroxy-cinnamic acid | 0.00 | 0.04 |  |  |  |  |
| 14 | 19.6 | *trans*-Ferulic acid | 0.86 | 1.17 | 0.44 |  |  |  |
| 15 | 19.9 | n-Alkanol C18 | 0.01 | 0.02 | 0.02 | 0.03 | 0.04 | 0.03 |
| 16 | 19.9 | ω-Hydroxyacid C14:0 | 0.00 | 0.00 | | 0.01 | 0.01 | |
| 17 | 20.4 | Acid C18:1 | 0.06 | 0.07 | 0.04 | 0.12 | 0.16 | 0.06 |
| 18 | 20.5 | Acid C18:2 | 0.02 | 0.02 |  | 0.05 | 0.05 |  |
| 19 | 20.6 | Ferulic acid TMS ester | 0.01 | 0.02 |  |  |  |  |
| 20 | 21.0 | Acid C18:1 TMS ester | 0.01 | 0.02 |  | 0.03 | 0.04 |  |
| 21 | 21.5 | Acid C19:0 (IS-2) | 10.27 | 13.34 | 4.89 |  |  |  |
| 22 | 21.9 | n-Alkanol C20 | 0.06 | 0.12 | 0.09 | 0.12 | 0.26 | 0.14 |
| 23 | 22.0 | ω-Hydroxyacid C16:0 | 2.84 | 1.61 | 0.51 | 5.47 | 3.60 | 0.77 |
| 24 | 22.2 | Acid C19:0 (IS-2) TMS ester | 0.23 | 0.33 | 0.03 |  |  |  |
| 25 | 22.6 | Acid C20:0 | 0.08 | 0.05 | 0.06 | 0.16 | 0.10 | 0.09 |
| 26 | 22.6 | $\alpha,\omega$-Diacid C16:0 | 2.53 | 1.64 | 1.12 | 4.89 | 3.66 | 1.68 |
| 27 | 22.9 | n-Alkanol C21 | 0.03 | 0.02 |  | 0.06 | 0.04 |  |
| 28 | 23.3 | $\alpha,\omega$-Diacid C16:0 α-TMS ester | 0.47 | 0.42 | 0.07 | 0.90 | 0.94 | 0.10 |
| 29 | 23.9 | ω-Hydroxyacid C18:1 | 11.30 | 6.11 | 8.80 | 21.79 | 13.61 | 13.17 |
| 30 | 24.0 | n-Alkanol C22 | 1.58 | 1.72 | 0.31 | 3.05 | 3.85 | 0.47 |
| 31 | 24.0 | ω-Hydroxyacid C18:0 | 0.75 | 0.23 | 0.05 | 1.44 | 0.52 | 0.07 |
| 32 | 24.5 | ω-Hydroxyacid C18:1 TMS ester | 1.40 | 4.87 | 0.23 | 2.69 | 10.87 | 0.35 |
| 33 | 24.6 | $\alpha,\omega$-Diacid C18:1 | 6.21 | 1.43 | 3.95 | 11.99 | 3.17 | 5.91 |
| 34 | 24.6 | Acid C22:0 | 0.43 | 0.54 | 0.69 | 0.83 | 1.20 | 1.04 |
| 35 | 24.7 | $\alpha,\omega$-Diacid C18:0 | 1.99 | 0.33 | 0.35 | 3.85 | 0.73 | 0.52 |
| 36 | 25.2 | $\alpha,\omega$-Diacid C18:1 α-TMS ester | 1.31 | 1.45 | 0.22 | 2.52 | 3.23 | 0.33 |
| 37 | 25.3 | $\alpha,\omega$-Diacid C18:0 α-TMS ester | 0.40 | 0.22 | 0.02 | 0.77 | 0.49 | 0.03 |
| 38 | 25.9 | ω-Hydroxyacid C20:1 | 0.04 | 0.05 | | 0.08 | 0.11 | |
| 39 | 26.0 | n-Alkanol C24 | 1.21 | 2.56 | 0.25 | 2.33 | 5.71 | 0.38 |
| 40 | 26.1 | ω-Hydroxyacid C20:0 | 0.65 | 0.45 | 0.53 | 1.26 | 1.00 | 0.79 |
| 41 | 26.1 | ω-Hydroxyacid C18:0, 9,10-dihydroxy | 0.83 | 0.83 | 3.83 | 1.60 | 1.86 | 5.73 |
| 42 | 26.3 | ω-Hydroxyacid C18:0, 9(10)-hydroxy-10(9)-methoxy | 0.02 | 0.02 | | 0.04 | 0.05 | |
| 43 | 26.4 | ω-Hydroxyacid C18:0, 9,10-epoxy | 0.71 | 1.19 | 5.72 | 1.36 | 2.64 | 8.56 |
| 44 | 26.6 | ω-Hydroxyacid C18:0, 9,10-dihydroxy TMS ester | 0.08 | 0.59 | | 0.16 | 1.33 | |
| 45 | 26.6 | ω-Hydroxyacid C20:0 TMS ester | 0.10 | 0.28 | | 0.20 | 0.63 | |
| 46 | 26.7 | Acid C24:0 | 0.46 | 0.92 | 0.39 | 0.89 | 2.04 | 0.58 |
| 47 | 26.7 | $\alpha,\omega$-Diacid C18:0, 9,10-dihydroxy | 0.55 | 0.31 | 4.94 | 1.07 | 0.69 | 7.40 |
| 48 | 26.8 | $\alpha,\omega$-Diacid C20:0 | 0.43 | 0.17 | 0.68 | 0.83 | 0.37 | 1.02 |
| 49 | 26.9 | n-Alkanol C25 | 0.07 | 0.75 |  | 0.13 | 1.67 |  |
| 50 | 27.0 | ω-Hydroxyacid C21:0 | 0.05 | 0.03 | | 0.09 | 0.06 | |
| 51 | 27.0 | ω-Hydroxyacid C21:0 TMS ester | 0.07 | 0.04 | | 0.13 | 0.08 | |
| 52 | 27.2 | $\alpha,\omega$-Diacid C18:0, 9,10-epoxy | 0.67 | 1.00 | 4.77 | 1.29 | 2.22 | 7.14 |
| 53 | 27.9 | n-Alkanol C26 | 0.29 | 1.69 | 0.05 | 0.57 | 3.77 | 0.07 |
| 54 | 28.0 | ω-Hydroxyacid C22:0 | 2.54 | 2.59 | 7.87 | 4.90 | 5.77 | 11.77 |
| 55 | 28.5 | ω-Hydroxyacid C22:0 TMS ester | 0.18 | 0.74 | 0.22 | 0.35 | 1.65 | 0.32 |
| 56 | 28.7 | Acid C26:0 | 0.32 | 0.45 |  | 0.61 | 1.00 |  |
| 57 | 28.8 | $\alpha,\omega$-Diacid C22:0 | 0.50 | 0.15 | 2.04 | 0.97 | 0.32 | 3.06 |
| 58 | 28.9 | ω-Hydroxyacid C23:0 | 0.03 | 0.02 | 0.03 | 0.05 | 0.05 | 0.04 |
| 59 | 29.2 | $\alpha,\omega$-Diacid C22:0 α-TMS ester | 0.05 | 0.05 | 0.13 | 0.09 | 0.10 | 0.19 |
| 60 | 29.7 | n-Alkanol C28 | 0.05 | 0.31 |  | 0.10 | 0.69 |  |
| 61 | 29.8 | ω-Hydroxyacid C24:0 | 1.21 | 2.03 | 1.20 | 2.34 | 4.51 | 1.80 |
| 62 | 30.2 | ω-Hydroxyacid C24:0 TMS ester | 0.01 | 0.03 | 0.04 | 0.02 | 0.06 | 0.07 |
| 63 | 30.4 | Acid C28:0 | 0.03 | 0.04 |  | 0.06 | 0.09 |  |
| 64 | 30.5 | $\alpha,\omega$-Diacid C24:0 | 0.23 | 0.15 | 0.15 | 0.45 | 0.33 | 0.23 |
| 65 | 30.6 | ω-Hydroxyacid C25:0 | 0.02 | 0.03 | | 0.04 | 0.06 | |
| 66 | 31.3 | ω-Hydroxyacid C26:0 | 0.30 | 0.41 | 0.03 | 0.58 | 0.91 | 0.05 |
| 67 | 31.9 | Sitosterol | 0.04 | 0.18 |  | 0.08 | 0.40 |  |
| 68 | 32.7 | ω-Hydroxyacid C28:0 | 0.05 | 0.10 | | 0.10 | 0.22 | |
| 69 | 36.4 | Ferulate of ω-hydroxyacid C16:0 | 0.10 | 0.05 | | 0.19 | 0.11 | |
| 70 | 38.3 | Ferulate of ω-hydroxyacid C18:1 | 0.17 | 0.05 | 0.24 | 0.34 | 0.11 | 0.35 |

Note: Peak numbers and retention time (RT) refer to GC-FID chromatogram elution order and are shown in Fig 6. IS – Internal standard. a - Carboxylic acid groups are in methyl ester form, and hydroxyl groups are in trimethylsilyl (TMS) ether form except where indicated. b - The area percentages are relative to the total area of identified compounds: 92% for (A) and 86% for (B). c - A relative response factor to standard (RRF) = 1 was used. Only identified monomers normally considered as part of suberin were considered. d - Extracted from raw data of Marques and Pereira (16) work. In this case, the data uses the same RRF=1.

**Table 4.** The mass percentage of the total identified components determines suberin monomeric composition by chemical families.

|  | % mass to suberin |  |  |
| --- | --- | --- | --- |
|  | A | B | *Q. suber*<sup>a</sup> |
| **Glycerol** | **14.9** | **10.9** | **17.3** |
| **Alkanols** | **6.4** | **16.0** | **1.1** |
| **Monoacids** | **2.9** | **4.8** | **1.8** |
| Saturated | 2.7 | 4.6 | 1.7 |
| Unsaturated | 0.2 | 0.2 | 0.1 |
| **$\alpha,\omega$-Diacids** | **29.8** | **16.5** | **27.7** |
| Saturated | 13.0 | 7.2 | 6.9 |
| Unsaturated | 14.5 | 6.4 | 6.2 |
| 9,10-Epoxy | 1.3 | 2.2 | 7.1 |
| 9,10-Dihydroxy | 1.1 | 0.7 | 7.4 |
| **ω-Hydroxyacids** | **44.9** | **50.2** | **43.5** |
| Saturated | 17.1 | 19.7 | 15.7 |
| Unsaturated | 24.6 | 24.6 | 13.5 |
| 9,10-Epoxy | 1.4 | 2.7 | 8.6 |
| 9,10-Dihydroxy | 1.8 | 3.2 | 5.7 |

<sup>a</sup> Extracted from raw data of Marques and Pereira (2019) work (16).

The methodology for suberin analysis, developed by Marques and Pereira (16), allowed for the simultaneous GC-FID analysis of glycerol and fatty components of suberin, as shown in the chromatogram in Fig 6, where glycerol appears as the highest peak (peak 1). This work employed a shorter GC column with higher polarity, facilitating the separation of the 70 identified components in less time than in the original study (Table 3).

Suberin analysis results rely highly on sample preparation and chemical analysis methodologies (16). For these reasons, Tables 3 and 4 show the suberin analysis results from *Q. suber*, where the same method was used but with the values refactored to an RRF=1 to the standards. Both *E. suberea* bark samples show a typical cork suberin composition pattern with an expected distribution of components where glycerol, ω- hydroxyacids and α,ω-diacids are the major contributors, with the suberin of the smallest tree (<u>A</u>) closer in composition to *Q. suber* cork. Suberin from the largest tree (<u>B</u>) reveals a composition with less glycerol, richer in alkanols, and poorer in α,ω-diacids. As observed for dichloromethane extractives, the difference can result from aging defense mechanisms.

Despite the high relative amount of unsaturated fatty acids (Table 4), the composition of *E. suberea* suberin shows a low content of mid-chain oxidation acids. This means there is a low level of epoxides and vic-diol functions in the ω-hydroxyacids and α,ω-diacids chains. The low level of epoxides can sometimes reflect an analytical artefact from unwanted hydrolysis during work-up, leading to an increased observed amount of vic-diol function, which is not applicable here. The low levels of epoxides and vic-diols in suberins are not abnormalities but specific features of that suberin biosynthetic pathway (55,56). This can also be observed in suberins from corks of other species, such as cork from Douglas-fir bark (40) and suberin from potato periderm (53).

The comparison of the suberin composition presented here with suberins from other species is often challenging because, in most cases, glycerol is either not quantified or is poorly analyzed. Additionally, in many analyses, standards are not employed, or sufficient care is not taken to prevent hydrolysis after methanolysis. However, when compared to bulk results from other suberins, *E. suberea* suberin displays an unusual characteristic: a significant ratio of its bifunctional fatty acids, *i.e.*, α,ω-diacids and ω- hydroxyacids, are disclosed as a trimethylsilyl ester (-COOTMS). This indicates that, originally *in situ*, these acid groups were free. These aliphatic acid monomers were linked to the suberin structure chain by only one of its two terminal functional groups, OH or COOH.

Table 5 illustrates the relative proportion of carboxylic acid functions analyzed as trimethylsilyl ester (TMS). Examining both suberin results, the bark from tree (<u>B)</u> shows 2 times more free acid end groups in α,ω- diacids and 3.6 times more free acid end groups in ω-hydroxyacids than the suberin from tree (<u>A)</u>. Compared to the TMS form of the standard, we can conclude that the hydrolysis of methyl esters was around 2-2.5% since the original free acid content in the standard was 0.05%. The hydrolysis of methyl esters was greater than that observed for *Q. suber*. This may have occurred during the methanolysis, considering that *E. suberea* bark appears to contain more structural water, as indicated by the FTIR spectrum showing a larger band around 1600 cm⁻¹. The presence of one free acid functionality in the bifunctionalized aliphatic suberinic acids suggests that these monomers were the terminal ends of the suberin structure. In the case of suberin from sample <u>B</u>, they represent one-third of the bifunctional monomers. *E. suberea* suberin seems to be a polymer with shorter aliphatic polyester chains than *Q. suber* suberin, and maybe some degradation of suberin structure can be associated with tree aging. These results follow the FTIR’s previous observations.

**Table 5.** The proportion of free fatty acids relative to the total of each family functional group in *E. suberea* suberin composition.

|  | % TMS form |  |  |
| --- | --- | --- | --- |
|  | A | B | *Q. suber* |
| C19:0 standard | 2.2 | 2.5 | 0.5 |
| **Acids** | **4.8** | **2.9** | **0.0** |
| **$\alpha,\omega$-Diacids** | **15.1** | **30.4** | **2.6** |
| **ω-Hydroxyacids** | **8.2** | **29.9** | **1.7** |

### 3.7. Lignin analysis

A summary of the pyrolysis analysis of the des-suberized barks is presented in Table 6, and the pyrograms are shown in Fig 7. The list of identified compounds and their relative percentages can be found in the supplementary material. The total carbohydrates are in smaller quantities and showed some degradation, as levoglucosan, the primary derivative of cellulose, was obtained in low amounts in samples <u>A</u> (0.7%) and <u>B</u> (1.2%). This may be due to the samples being des-suberized before this analysis, which could lead to some alterations in the cellulose. However, the cellulose content is not as high as other cork-containing barks. The value found in eucalypt wood ranges from 22.7% to 27.5% (57), while in the bark from stumps, it was 31.9% (58). For other barks containing cork, the values obtained from this analysis were 40% in Douglas-fir cork (50) and 42.5% total in *Parinari curatellifolia*, where these last samples were also des- suberized previous study (59).

**Fig 7.**
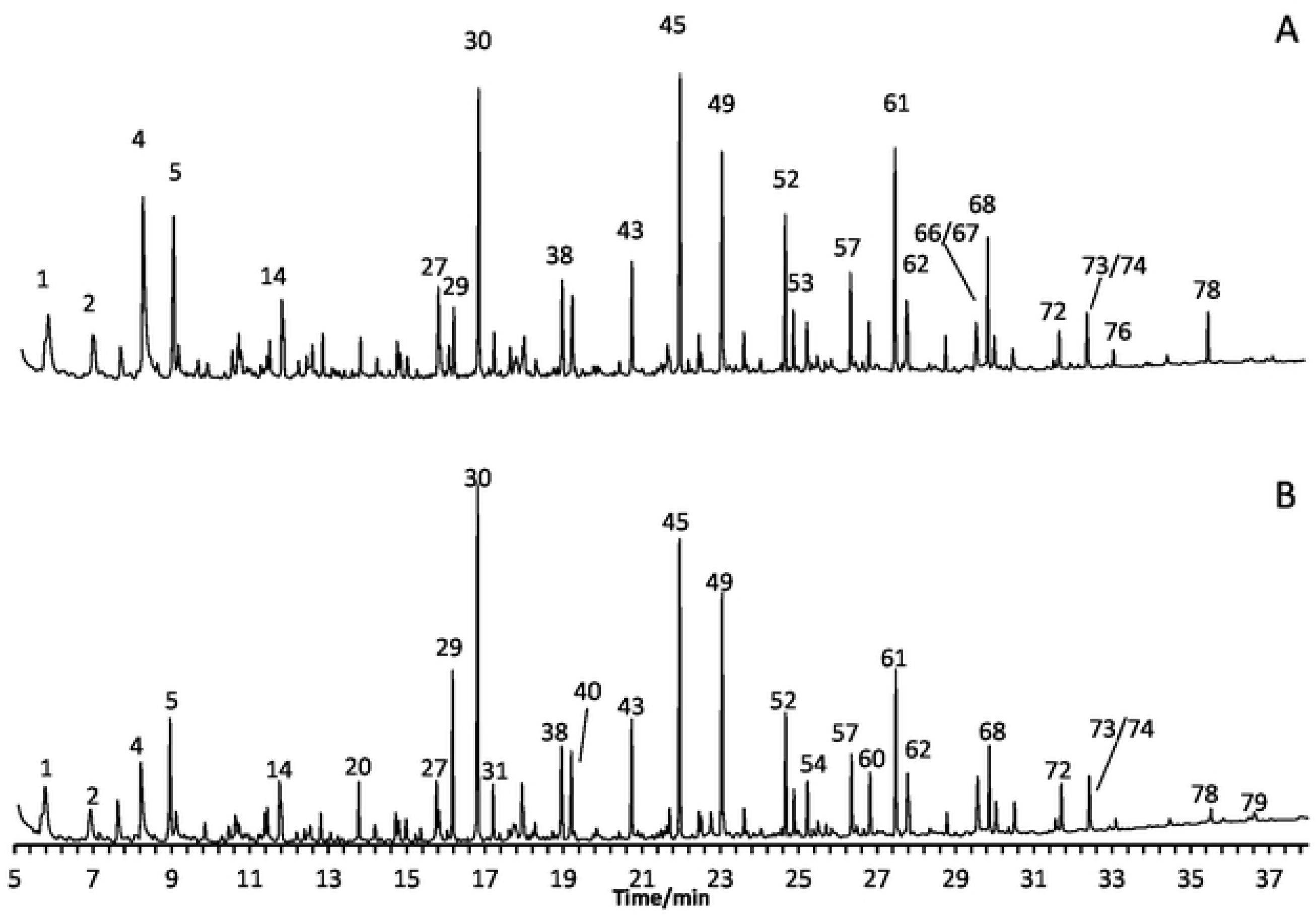
Pyrograms of the barks <u>A</u> and <u>B</u> off. *suberea.* Legend: 1) acetone & 2-oxo-propanal; 2) 2,3-butanedione; 4) acetic acid; 5) 1-hydroxy-2-propanone (acetol); 14) furfural; 20) 2-hydroxy-2-cyclopenten-1-one; 27) 3-methyl-1,2-cyclopentanedione & 2,3-dimethyl-2-cyclopenten-1-one; 29)phenol; 30) guaiacol; 31) o-cresol; 38) 4-methylguaiacol; 40) CH_2_=COH-CHOH-CO-CH,-CH_3_; 43) 4-ethylguaiacol; 45) 4-viny1guaiacol; 49) syringol; 52) *trans* isoeugenol; 53) 4-methylsyringol; 54) vanillin; 57) 4-ethylsyringol; 60) acetoguaiacone; 61) 4-vinylsyringol; 62) guaiacylacetone;66) 1,6-anhydro-β-D-glucopyranose; 67) 4-propinylsyringol; 68) *trans* 4-propenylsyringol; 72) acetosyringone; 73) syringylacetone; 74) *trans* coniferaldehyde; 76) ferulic acid methyl ester; 78) 17-methyl octadecanoic acid methyl ester; 79) *trans* sinapaldehyde.

**Table 6.** Summary of the derived compounds of lignin and carbohydrates attained by pyrolysis of des-suberized bark samples from *E. suberea*.

| A sum of identified compounds (as % total area) |  | <u>A</u> | <u>B</u> |
| --- | --- | --- | --- |
| Total carbohydrates |  | 34.4 | 31.9 |
| Total lignin |  | 46.9 | 57.2 |
| others |  | 3.0 | 3.8 |
| Sum of lignin derivatives (% of total lignin) |  |  |  |
| S |  | 36.8 | 33.7 |
| G |  | 55.5 | 52.9 |
| H |  | 7.8 | 13.4 |
| S/G ratio |  | 0.66 | 0.64 |
| C/L ratio |  | 0.73 | 0.56 |
| H:G:S relation |  | 1:7:5 | 1:4:3 |

According to Py-GC-MS, both des-suberized barks of *E. suberea* are rich in lignin, especially bark <u>B</u> (57.2%) compared to bark <u>A</u> (46.9%), containing a GS-type lignin with a S/G ratio of 0.66 and a monomeric H:G:S ratio of 1:7:5 for sample <u>A</u>, while sample <u>B</u> achieved S/G values of 0.64 and H:G:S of 1:4:3 (Table 6). These results are in accordance with FTIR analysis. Lignin/carbohydrate ratios registered here by Py-GC-MS are higher than those recorded in the summative chemical analysis due to the partial hydrolysis of hemicelluloses that occur during the methanolysis of suberin and which is proven by the unidentified sugars present in the methanolysis extract of suberin. The bark of *E. suberea* has a different lignin composition, with a higher percentage of S-units, when compared with other important corky-bark species, such as *Q. suber*, *Q. cerris*, *Betula pendula,* and *P. menziesii,* where G-units prevail over S-units (17,50,60) and the S/G values are under 0.14. Other barks, such as the bark of *Parinari curatellifolia*, with an S/G ratio of 0.32 (60), or *B. recurvata*, with 0.40 (61), present higher S/G ratios. The composition of lignin appears variable and dependent on the origin of the bark, with no apparent relationship with the suberin content or the plant group (gymnosperm or angiosperm). However, these GS-lignin characteristics are quite distinct from the cork of *Quercus suber*, which has more guaiacyl units and fewer syringyl units, resulting in an S/G ratio ranging from 0.029 to 0.10 (17,62,63). The few studies carried out to date, including isolation, composition, and structure of cork lignin, have revealed that, regardless of the species of cork origin, cork lignin is a Guaiacyl-type lignin (17,50,60). Studies demonstrated a composition of ca. 95% of guaiacyl units with minor amounts of syringyl monomers and incorporation of a variable amount of ferulic acid as a construction guaiacyl-type monomer in its structure that differentiates this lignin from those present in woods (18). Ferulic acid is always present in cork analyses, also identified by analytical pyrolysis in its methyl ester form (peak 76); being a builder of interconnection bridges between the different aromatic and aliphatic domains, plays an important role in the construction of the cell wall, and has been one of the missing links in analysis methodologies.

## Conclusions

In conclusion, this work presents, for the first time, an anatomical and chemical characterization of *E. suberea* bark. The outer bark of *E. suberea* is composed mainly of cork. It can be ground into coarse granules directly without needing any prior physical separation, which is important in enhancing possible uses, namely in manufacturing agglomerates and composites. Microscopic observations reveal that the bark contains cells typical of cork tissue. The chemical characterization indicates: *i)* a lipophilic extractive composition made up of a mixture of fatty components, n-alkanols, acids, monoglycerides, and particularly ferulates of n-alkanols, with ferulates and α,ω-bifunctional acids accounting for 27% and 25% of this extract, *ii)* a suberin composition pattern with a predictable distribution of components, where glycerol, ω- hydroxy acids, and α,ω-diacids are the major compounds, *iii)* a suberin structure with shorter aliphatic polyester chains compared to *Q. suber* suberin and *iv)* a lignin monomer composition with an S/G ratio between 0.64 and 0.66, prevailing guaiacyl units but in less percentage compared to cork of *Q. suber*.

The results of this study indicate that the bark of *E. suberea* is a raw material with the necessary characteristics to be used as an interesting alternative for producing products derived from cork in response to the increasing scarcity of this material as well as being a species that can present a better adaptation to climate change due. Nevertheless, for using *E. suberea* as a wood and cork source, more studies are necessary concerning forestry exploitation techniques, wood physical and chemical studies, wood pulp production, and cork material physical behavior.

## Conceptualization

A.V.M and JG; methodology: A.L., A.V.M, and JG; validation: A.L., A.V.M and JG; formal analysis: A.L., A.V.M, and JG; investigation: T.Q, H.P, A.L., A.V.M, and JG; data curation: T.Q, H.P, A.L., A.V.M, and JG; writing-original draft preparation: A.V.M; writing-review and editing: T.Q, H.P, A.L., A.V.M and JG. All authors have read and agreed to the published version of the manuscript.

## Acknowledgments

This research was funded by the Portuguese Foundation for Science and Technology (FCT) through its support of the Forest Research Centre (UIDB/00239/2020) and the Project CLEAN FOREST (PCIF-GVB-0167- 2018). Ana Lourenço acknowledges a research contract (DL 57/2016/CP1382/CT0007, https://doi.org/10.54499/DL57/2016/CP1382/CT0007), and Helena Patrício by a doctoral fellowship (2023.04037.BD) which FCT also funds. The authors thank Carolina Godinho for her help with the chemical analysis. They also thank INIAV for allowing the collection of the raw material plantation in Escaroupim National Forest.

